# An organismal view of newborn cell dynamics in mammalian aging

**DOI:** 10.64898/2026.05.26.728032

**Authors:** Ziyu Lu, Wei Zhou, Junyue Cao

**Author notes:** Senior author (Wei Zhou, Junyue Cao). Correspondence (W.Z.), (J.C.).

## Abstract

Aging remodels tissue composition, yet static single-cell atlases cannot show which renewal processes drive this change across the organism. To address this gap, we applied *TrackerSci* to 21 mouse tissues at young, adult, and aged stages in both sexes, linking *in vivo* EdU labeling to single-nucleus chromatin accessibility. This produced approximately 3 million profiles of newborn cells matched to global cell populations from the same tissues and ages. Newborn-cell abundance varied widely across tissues and generally declined with age. Beyond abundance, the cell-type composition of newborn cells differed from that of the tissue as a whole. Reflecting this difference, they showed distinct chromatin states, including known renewal compartments and candidate progenitor or proliferation-active states. With age, they also shifted along inferred differentiation trajectories, with stalled-differentiation patterns emerging in multiple lineages. At the population level, age-related changes in newborn-cell abundance partially explained the broader shifts in cell-type composition during aging. Renewal-rate analysis further showed that age-related cell-type declines were driven by reduced replacement, while expansions reflected altered cell-state transitions. Together, these findings show how newborn cell dynamics across the whole organism reveal the routes by which aging remodels cellular composition.

## Introduction

Aging remodels tissues not only by changing which cells are present, but by altering the routes through which cell populations are produced, maintained, and lost (*1*, *2*). Cellular composition is a demographic endpoint. It reflects cell production, differentiation, persistence, redistribution, and clearance, but it does not reveal which of these processes has changed (*3–7*). This distinction is central to aging biology, where stem-cell exhaustion and reduced renewal capacity (*2*, *8*) have been linked to dysfunction in hematopoietic (*9*, *10*), muscle(*11*, *12*), intestinal (*13*, *14*), and neural systems (*15*, *16*). Yet even where stem-cell aging has been documented, static cellular composition alone cannot determine which process drove the observed change.

Measuring these dynamics across the organism has remained challenging. Classical *in vivo* labeling, proliferation, and lineage studies define cellular dynamics with mechanistic depth, but typically in selected lineages or reporter-defined populations (*6*, *17–22*). Single-cell transcriptomic and chromatin atlases, in contrast, have transformed aging biology by mapping cell types and states across tissues, organs, and lifespan stages (*23–25*). These atlases define what is present at sampling, but not the cell production and differentiation processes that gave rise to those states. A cell-type-specific, organism-wide measurement of newborn cell production is therefore needed to connect aging-associated composition to the dynamics that generate it.

We previously developed *TrackerSci* (*26*), a single-cell combinatorial-indexing method that couples EdU labeling of newly synthesized DNA with single-cell molecular profiling to study progenitor dynamics in the mammalian brain. To extend this framework organism-wide, we applied *TrackerSci* across 21 mouse tissues at three lifespan stages (1, 5, and 21 months) and in both sexes after a five-day EdU-labeling window, mapping newborn cell populations across the body. The resulting dataset comprises ∼3 million single-nucleus chromatin accessibility profiles of newborn cells, allowing us to define the *in vivo* proliferative landscape across cell types, tissues, and ages. We used this dataset to ask how newborn cell dynamics contribute to cellular remodeling during aging.

Our study revealed how newborn cells contribute to aging-associated tissue remodeling. Specifically, the abundance of these cells varied widely across tissues and generally declined with age, and they frequently showed chromatin states distinct from matched global populations. With age, they also shifted along inferred differentiation trajectories, with stalled-differentiation patterns emerging in selected lineages. Across cell types, age-related changes in newborn-cell abundance were positively associated with matched global population changes, but explained only part of compositional remodeling. Together, these findings show that organism-wide newborn cell dynamics reveal how production, maintenance, and loss shape cellular composition during aging.

## Results

### Global profiling of newborn cell dynamics across the mouse lifespan

To map newborn cell populations and their dynamics during aging, we applied *TrackerSci*(*26*) to profile EdU-labeled cells across 21 mouse tissues at young (1 month), adult (5 months), and aged (21 months) stages of both sexes (**Figure 1A, Table S1**). Briefly, mice were labeled with EdU for five consecutive days to mark *in vivo* cellular proliferation. Nuclei were then extracted from each tissue, followed by click chemistry–based fluorophore labeling(*18*) and fluorescence-activated cell sorting (FACS) to enrich the EdU+ newborn population. Sorted nuclei were then profiled by *EasySci-ATAC*(*27*) to generate single-nucleus chromatin accessibility libraries (**Figure 1A**). FACS-based quantification showed that cell genesis rates varied substantially across tissues in adult mice, with the highest activity in immune organs such as bone marrow (60.5% EdU+ cells), thymus (58.6%), and spleen (16.0%) (**Figure 1B and S1A, Table S1**). Skin (23.9%) and the gastrointestinal tract (stomach, 11.6%; intestine, 22.5%; cecum, 27.5%; colon, 17.1%) also showed substantial turnover. In contrast, most solid organs (kidney, liver, lung, heart, and muscle) showed low EdU+ fractions (*e.g.,* heart, 1.98%; muscle, 2.67%), indicative of limited renewal (**Figure 1B and S1A**).

**Figure 1.**
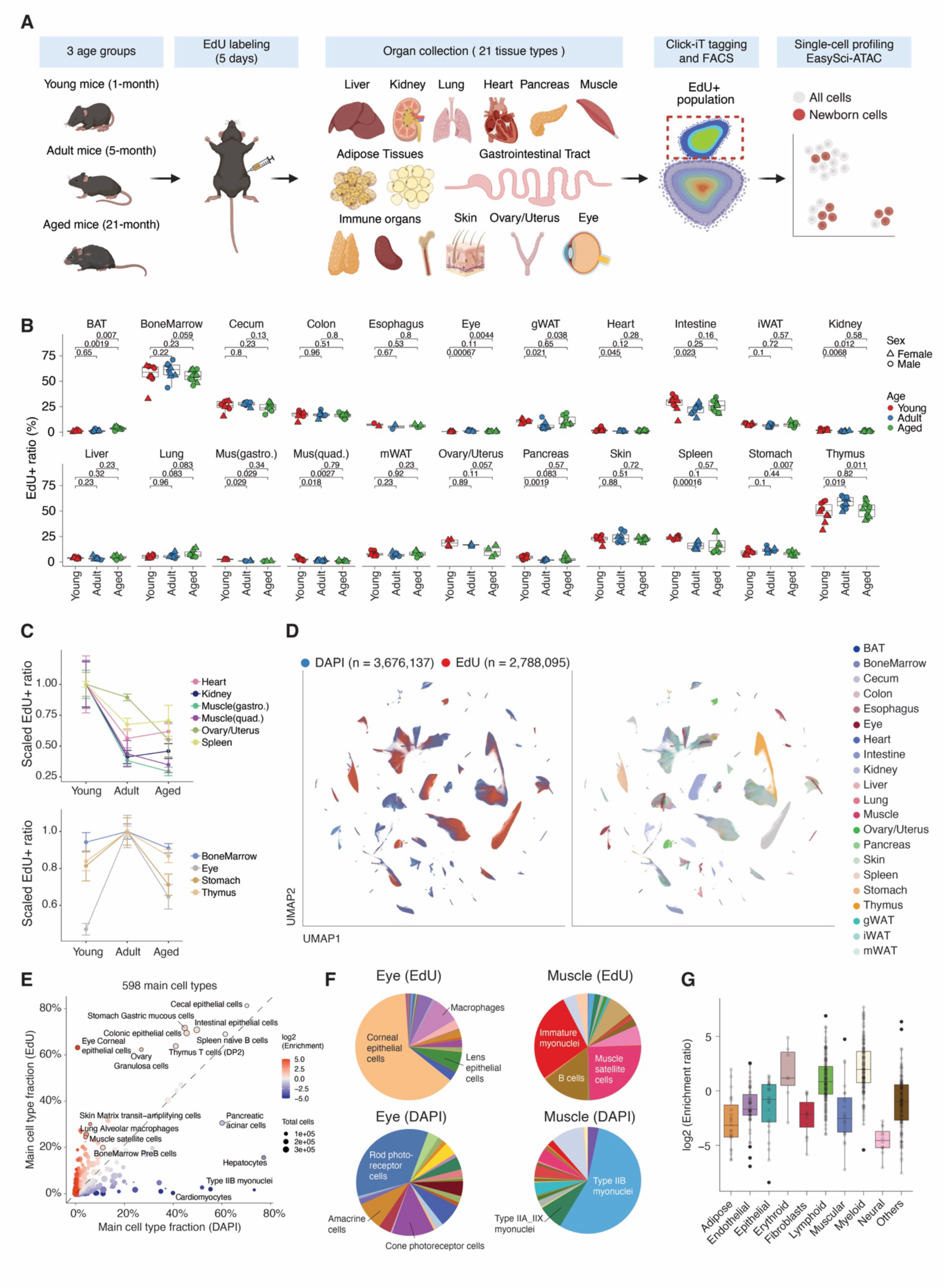
Global profiling of newborn cell dynamics across the mouse lifespan. **(A)** Schematic of the experimental workflow. **(B)** Box plot showing the fraction of EdU+ nuclei measured by FACS in each of the 21 tissues, stratified by age. Each point is one biological replicate. Point shape denotes sex. Pairwise comparisons between age groups use the two-sided Wilcoxon rank-sum test, with P-values labelled. **(C)** Line plot showing age trajectories of the mean EdU+ fraction for representative tissues, normalized to each tissue’s maximum across the three time points. Top: a monotonic age-related decline. Bottom: a non-monotonic trajectory peaking at the 5-month adult stage. Points show mean ± SEM across biological replicates. **(D)** UMAP embedding of single-nucleus chromatin accessibility profiles from the joint clustering of EdU+ (newborn) and DAPI (global) populations. Cells are colored by sorted population (left) and by tissue of origin (right). For the DAPI cells, we subsampled up to 5,000 cells per cell type before the co-embedding. **(E)** Cell-type composition of EdU+ and DAPI populations across main cell types in 1-month-old mice. Each point represents a tissue-specific cell type, showing its fraction within DAPI (x-axis) versus EdU+ (y-axis) populations. Proportions were quantified within each respective tissue. **(F)** Pie charts of cell-type composition in EdU+ and DAPI populations for two representative tissues (eye and muscle). The top-represented cell types within each population are labeled. **(G)** Box plots of lineage labeling enrichment across the whole organism in 1-month-old mice. Each point represents a tissue-specific cell type. Enrichment is defined as the ratio of a cell type’s fraction in the EdU+ population to its fraction in the DAPI population.

We next examined how overall proliferation changes with age across the body. In the majority of tissues, including the heart, kidney, and skeletal muscle, the EdU+ fraction declined continuously across the three time points, reflecting a progressive loss of cellular renewal during aging (**Figures 1B-C**). A subset of tissues, including bone marrow, eye, stomach, and thymus, instead peaked at the adult stage before falling at the aged stage (**Figure 1C**). Across both trajectories, EdU+ fractions ultimately fell by the aged stage, indicating that age-associated loss of cellular renewal is broadly conserved. The specific trajectory of turnover, however, is tissue-dependent, likely reflecting shifts in the contributions of distinct proliferating cell types within each tissue.

To resolve the identity of these newly generated cells, we jointly clustered the global (DAPI+) cells profiled in our previous study(*25*) and newborn (EdU+) populations, recovering 598 organ-specific main cell types (**Figure 1D and Figure S2**). The global and newborn populations showed markedly different cell-type compositions (**Figure 1E**). For example, some abundant cell types (pancreatic acinar cells, hepatocytes, and muscle myonuclei) were strongly underrepresented in EdU+ relative to DAPI, consistent with their quiescent state. Conversely, several rare cell types (corneal epithelial cells and immature myonuclei) occupied a disproportionately large fraction of the EdU+ population, confirming the specificity of *TrackerSci* to enrich actively renewing pools (**Figure 1F**). When aggregated by lineage, lymphoid, myeloid, and erythroid cells were most enriched in the newborn fraction, whereas neural cells showed the lowest turnover (**Figure 1G**). This lineage-level pattern is consistent with prior estimates in humans (*1*), and prompted us to investigate whether newborn cells are molecularly distinct from their homeostatic counterparts.

### Cellular state differences between newborn and global populations

We next examined whether, within each cell type, the newborn cells are simply a random subset of all cells or instead exhibit unique chromatin accessibility landscapes. To quantify this, we computed the energy distance(*28*) (a non-parametric measure of distributional difference) between EdU+ and DAPI cells for each main cell type, and assessed significance by permutation testing (**Figure 2A, Methods**). Of the 330 main cell types with sufficient cells in both populations for testing, 248 (75.2%) showed a significant EdU+ vs DAPI distribution shift (**Figure 2B**, FDR < 0.001). These results indicate that newborn cells are not a homogeneous reflection of their parent population. Instead, they form molecularly distinct subsets, likely corresponding to localized progenitors or highly proliferative pools.

**Figure 2.**
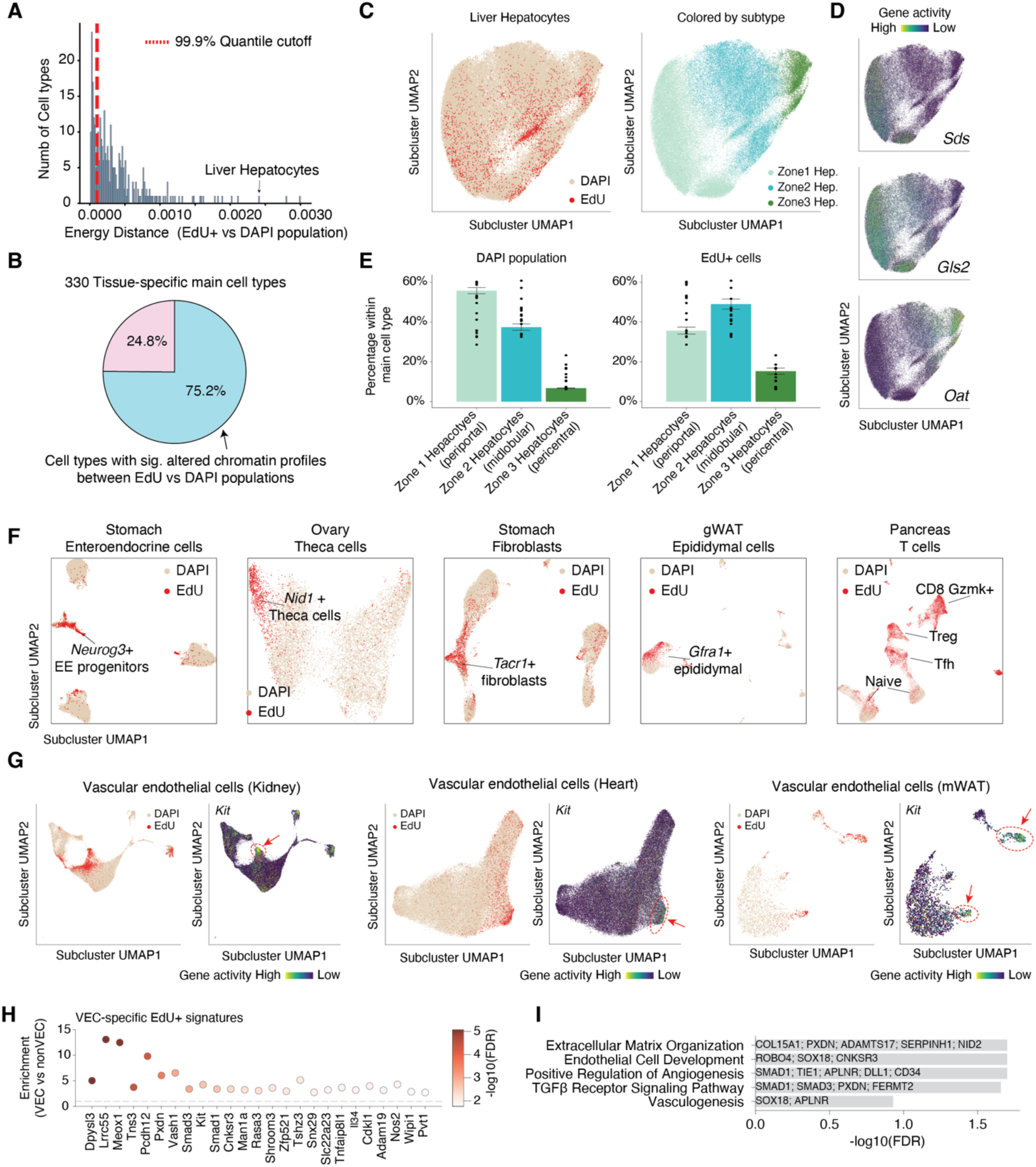
Cell state differences between newborn and global populations within cell types. **(A)** Histogram of the observed energy distance distribution across all cell types. The red dashed line denotes the significance threshold (FDR = 0.001) derived from a null distribution. **(B)** Pie chart showing the percentage of tissue-specific main cell types with a significant shift in their chromatin accessibility between EdU and DAPI population. **(C)** Sub-cluster UMAP of liver hepatocytes, colored by EdU/DAPI (left) or by zonation subtype (right). **(D)** Sub-cluster UMAP of liver hepatocytes, colored by zonation markers *Sds*, *Gls2* and *Oat* (*31*). **(E)** Bar plots with SEM show the mean of percentage assigned to each zonation subtype. Dots represent individual biological replicates. **(F)** Sub-cluster UMAPs of example main cell types in which EdU+ cells localize to a discrete progenitor or proliferating sub-state. Cells are colored by the EdU/DAPI label. Treg: Regulatory T cells; Tfh: T follicular helper cells. **(G)** Sub-cluster UMAP colored by EdU/DAPI and gene accessibilities of *Kit* in vascular endothelial cells (VECs) from kidney, heart, and mWAT. **(H)** Top VEC-specific EdU+ signature genes. X-axis: genes ranked by Fisher’s exact enrichment p-value, top 25 shown; y-axis: enrichment ratio, defined by fraction of VEC cell types in which the gene is significantly up in EdU+ vs DAPI, divided by the matching fraction across all non-VEC main cell types. **(I)** Barplot showing top enriched gene ontology terms for the VEC-specific EdU+ signature in (H).

A representative example is liver hepatocyte zonation. Hepatocytes are spatially organized into three zones along the liver lobule: zone 1 (periportal), zone 2 (midlobular), and zone 3 (pericentral).(*29*, *30*) Subclustering of our liver hepatocytes recovered all three zones (Figure 2C), confirmed by canonical zonation markers(*31*, *32*) (**Figure 2D**). The global DAPI population was dominated by zone-1 hepatocytes, whereas EdU+ cells were enriched in zone 2 (**Figure 2E**), consistent with recent lineage-tracing reports that pinpoint zone 2 as the source of homeostatic hepatocyte renewal (*31*, *32*).

Beyond confirming known cases such as the liver zonation axis, our approach also enabled the *de novo* identification of progenitors or highly proliferative subpopulations across diverse tissues. These included *Neurog3*+ enteroendocrine progenitors (*33*) and *Tacr1*+ fibroblasts (stomach), *Nid1*+ theca cells (ovary) (*34*), *Gfra1*+ epididymal cells (gonadal white adipose tissue, gWAT) (*35*, *36*), and proliferation-active T cell states including CD8 *Gzmk*+ T cells, regulatory T cells, and follicular helper T cells (pancreas) (**Figure 2F and Figure S3**). These examples reveal a previously unappreciated diversity of cell-type-specific progenitor or actively cycling populations across mammalian tissues, many of which were not resolvable by canonical markers alone.

The organism-wide scope of our dataset further enabled the identification of conserved progenitor states shared across tissues for broadly distributed cell types. For example, in endothelial cells (EC) from the kidney, heart, and mesenteric white adipose tissue (mWAT), we identified an EdU-enriched population with high accessibility of *Kit*, a known endothelial progenitor gene(*37*, *38*) (**Figure 2G**). To confirm this conserved EC progenitor identity, we performed differential gene-accessibility analysis (Wilcoxon test, EdU+ vs DAPI) within each main cell type, and searched for genes whose EdU+ enrichment was specific to endothelial cells. This identified several candidate EC progenitor genes, including the known endothelial regulator *Meox1*(*39*) (**Figure 2H, Table S3**). Pathway enrichment analysis showed that these genes were significantly enriched for endothelial biology terms, such as extracellular matrix organization and positive regulation of angiogenesis (**Figure 2I**), supporting their identity as endothelial progenitors.

### Cellular differentiation trajectories change with aging

We next interrogated cell differentiation processes, leveraging EdU labeling not only to tag proliferating progenitor cells but also to track the differentiation of their progeny over time. We focused on ten representative trajectories spanning multiple tissues, including immune cell genesis in the bone marrow and thymus, epithelial differentiation in the intestine and colon, epidermal differentiation in the skin, and an aging-associated trajectory in the kidney that connects parietal epithelial cells (PECs) to the PTS3T2 state (**Figure 3A**). As expected, when overlaying EdU+ cells onto the global cell population, newborn cells were enriched in early differentiation states, whereas the global population spanned the entire trajectory. Building on this, we computed pseudotime for each trajectory (**Figure 3A**). We confirmed directionality using canonical stem-cell markers (**Figure S4A and S4B**, *e.g., Cd34*+ HSPCs (*40*) in the bone marrow and *Lgr5*+ epithelial stem cells in the intestine (*6*)).

**Figure 3.**
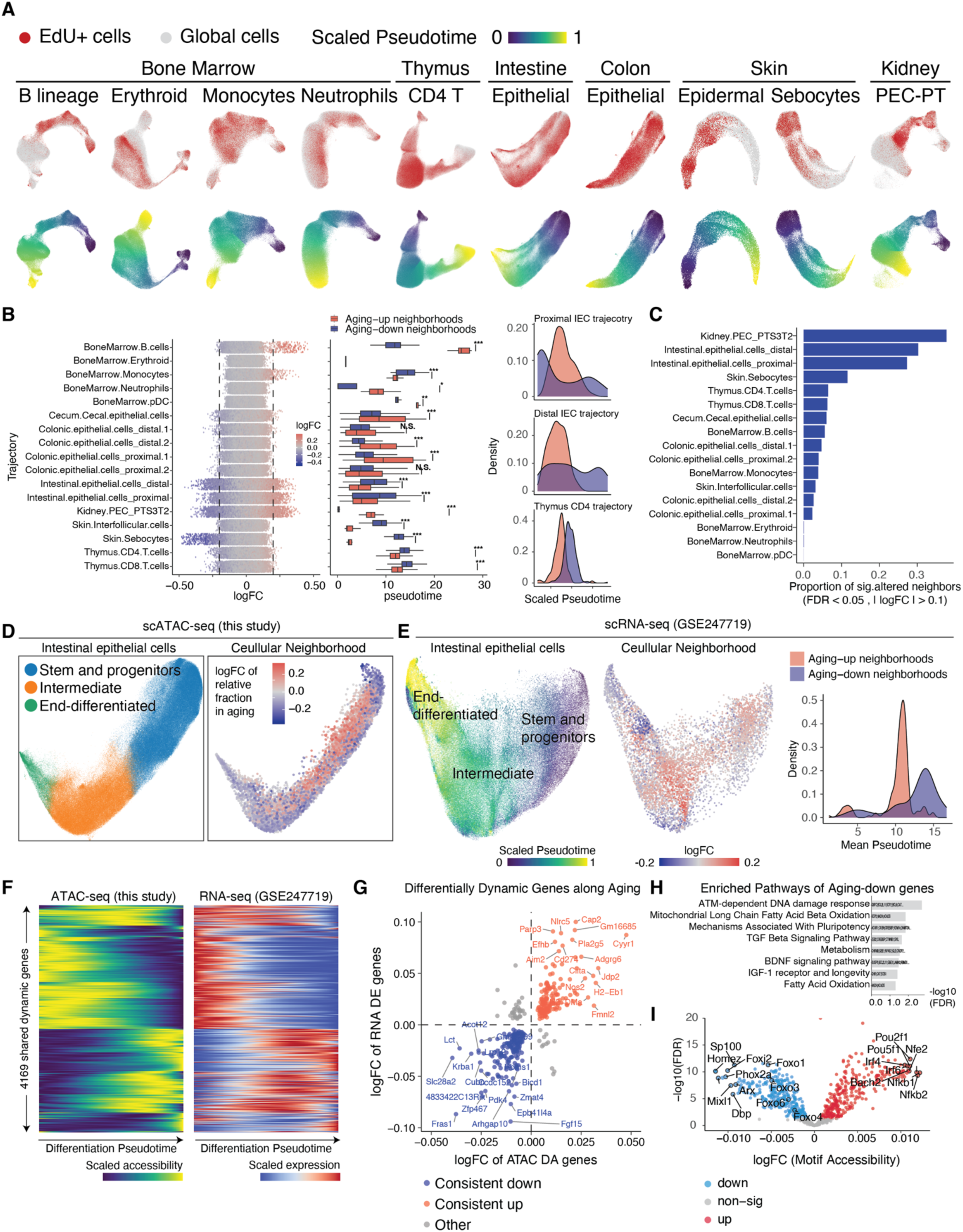
Aging affects cellular differentiation across multiple tissue trajectories. **(A)** UMAP plot of ten representative differentiation trajectories across five tissues. Top row: EdU+ newborn cells (red) overlaid on the global cell population (gray). Bottom row: the same embeddings colored by Slingshot pseudotime, scaled to [0, 1]. **(B)** Differential abundance testing of EdU+ cells along aging using Milo. Left: per-neighborhood log2 fold change (Aged vs. Young) for each trajectory. Middle: boxplot of pseudotime-distribution of aging-associated neighbors. Right: density of aging-enriched (red) and aging-depleted (blue) neighborhoods for three example trajectories. **(C)** Proportion of significantly altered neighborhoods (FDR < 0.05, |log2FC| > 0.1) per trajectory. **(D)** Cellular neighborhood analysis of the proximal intestinal epithelial trajectory in scATAC-seq (this study). Left: UMAP colored by cell state. Right: same UMAP with neighborhoods colored by log2 fold change of relative cell-fraction in aging (red = aging-enriched, blue = aging-depleted). **(E)** Validation in scRNA-seq from an independent dataset. Left: UMAP of intestinal epithelial cells colored by scaled pseudotime. Middle: same UMAP colored by aging log2 fold change at the cellular-neighborhood level. Right: density of aging-enriched and aging-depleted neighborhoods along mean pseudotime, recapitulating the stalled-differentiation pattern observed in (D). **(F)** Heatmaps of 4,169 dynamic genes shared between ATAC-seq (left, this study) and RNA-seq (right, GSE247719) along the proximal intestinal epithelial differentiation pseudotime. Rows are genes ordered by peak position along pseudotime; color encodes scaled accessibility (left) or scaled expression (right). **(G)** Scatter plot of dynamic genes affected by aging, comparing log2 fold change in ATAC-seq gene activity (x-axis) against log2 fold change in RNA-seq expression (y-axis). Genes are colored based on concordant directionality across the two modalities. **(H)** Pathway enrichment of aging-down dynamic genes. **(I)** Volcano plot of transcription-factor motif accessibility in the intestinal epithelial trajectory.

To assess whether and how aging perturbs these differentiation trajectories, we applied the differential abundance algorithm *Milo*(*41*) to EdU+ cells, identifying newborn-cell neighborhoods that are significantly altered with aging (**Figure 3B, left**). Across the selected trajectories analyzed, 14 of the 17 exhibited significant shifts in pseudotime between aging-up and aging-down neighborhoods. A recurring pattern in aged samples was the accumulation of cells at intermediate stages of differentiation, coupled with a depletion of early progenitor or late mature stages (**Figure 3B, middle**). These shifts indicate that aging slows down progenitors and also delays differentiation before cells reach maturity. This stalling pattern was conserved across multiple lineages, including the intestinal epithelium and thymic T-cells (**Figure 3B, right**), indicating it is a broadly shared feature of aging across the body.

Although the altered differentiation pattern was broadly shared, trajectories differed in the magnitude of aging effects. Ranking them by the proportion of neighborhoods significantly changed with aging (**Figure 3C**), we found that the PEC-to-PTS3T2 trajectory was most strongly affected. This trajectory connects parietal epithelial cells (PECs), normally non-proliferative cells of the renal corpuscle(*42*), to a proximal tubule subtype (PTS3T2) (*43*), which we previously reported expands specifically in aged female kidneys (*25*). Sub-clustering of the PEC compartment identified the cellular source of the aging-associated expansions of PTS3T2. We resolved three distinct subpopulations, where a novel, actively proliferating Ror2+/Dock2+ population emerged with aging (**Figure S4G**) unique to females. This suggests a potential newborn-state source associated with the sex-biased expansion of PTS3T2 cells (**Figure S4H**) (*25*).

Turning to a canonical differentiation trajectory, we next examined the epithelial trajectory of the proximal intestine in depth (**Figure 3D**). This trajectory is a representative case of the stalled-differentiation pattern: with aging, newborn cells accumulate at intermediate stages, with depletion at both the early progenitor and the late differentiation ends. To validate this pattern in an orthogonal data modality, we examined the same cell type in published scRNA-seq data(*24*). We integrated ATAC and RNA modalities to infer pseudotime for the scRNA-seq cells (**Figure 3E, left**), and recovered the same delayed progression in this independent dataset (**Figure 3E, middle and right**).

To investigate the molecular basis of this stalling, we first defined dynamic genes whose accessibility or expression changes along pseudotime, and then asked which of these were significantly altered in aging. We identified 4,169 dynamic genes with consistent patterns across ATAC and RNA (**Figure 3F**). Among them, 207 were significantly altered with consistent direction of change in aging in both modalities (**Figure 3G, Table S4**). Pathway enrichment of the aging-down genes identified pluripotency mechanisms, TGF-β signaling (*44*), mitochondrial β-oxidation (*45*), IGF-1 signaling (*46*), and DNA damage response (*47*) (**Figure 3H**). This convergent loss of stemness, metabolic, and differentiation programs is consistent with progenitors lacking both the capacity to maintain themselves and the support needed to complete differentiation. To identify TFs with aging-altered activity, we quantified motif accessibility changes from the ATAC-seq data (**Figure 3I**). Foxo motifs (*Foxo1, Foxo3, Foxo4*) showed decreased accessibility with aging, whereas inflammatory motifs (*Irf3, Irf4, NfkB*) showed increased accessibility, recapitulating the loss of Foxo program in IEC differentiations (*48–50*) and inflammatory activation in aging (*51–53*).

### Newborn cell changes in aging and their association with the global cell population

Having characterized how aging altered differentiation within individual trajectories, we next asked whether aging also changes the overall abundance of newborn cells across cell types and organs. For each main cell type in each sample, we calculated a “newborn ratio,” defined as the fraction of that cell type within the newborn pool, multiplied by the EdU labeling rate for each sample. We then applied regression analysis to identify cell types whose newborn ratio changed significantly with age within each organ, considering age-associated changes that differ between sexes (**Methods**). Because aging-associated changes may differ substantially between early and late life, we performed this analysis separately for the young-to-adult and adult-to-aged transitions. In total, we identified 227 organ-specific cell types with significantly altered newborn ratios across the two transitions (186 in the young-to-adult transition, 76 in the adult-to-aged transition, and 35 in both), with 37 showing significant age-by-sex interactions (FDR < 0.05; **Figure 4A, Table S5 and S6**).

**Figure 4.**
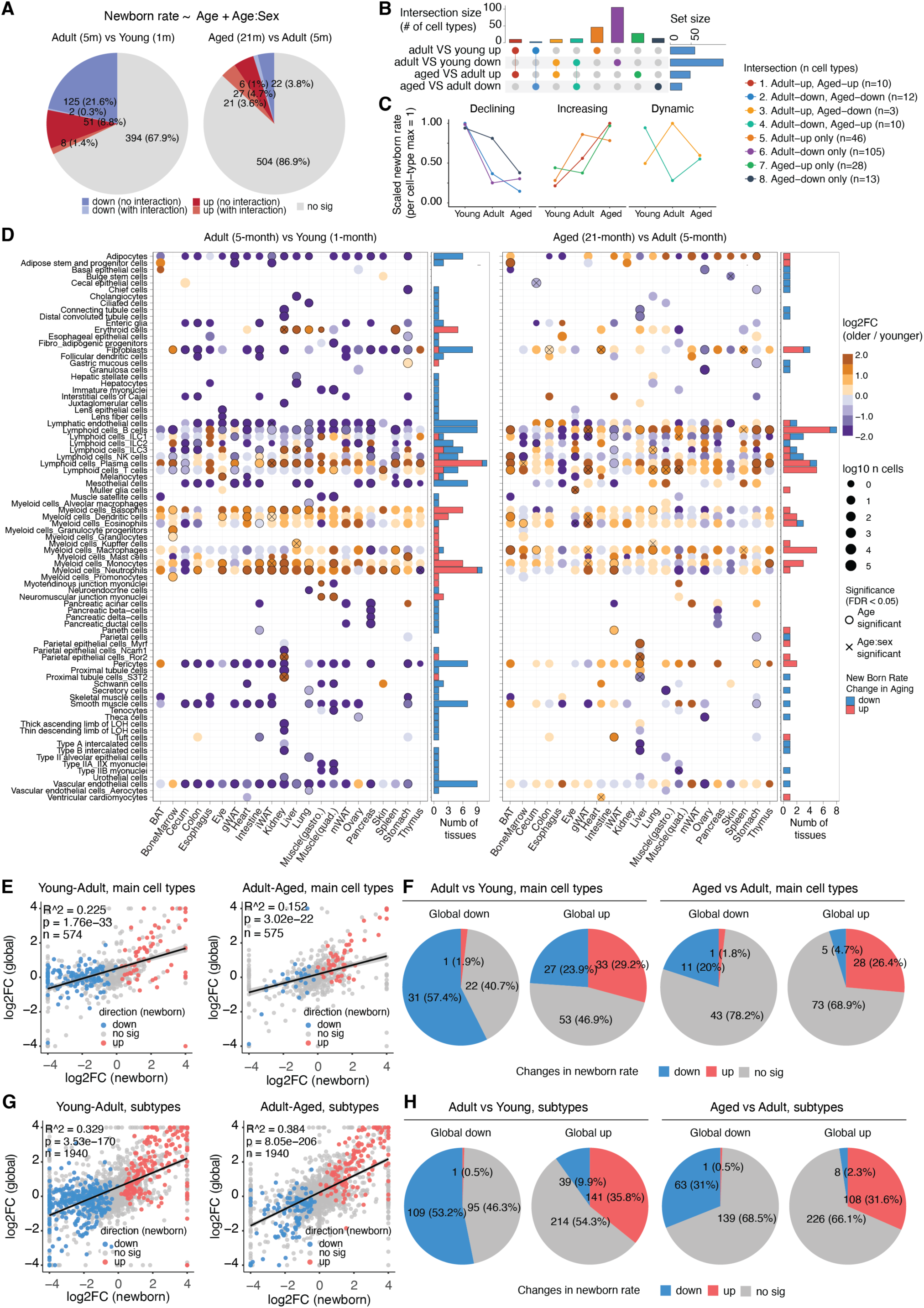
Aging-associated changes in the newborn cell population and their relationship to global cell-population shifts. **(A)** Pie charts summarizing organ-specific cell types whose newborn ratio changes significantly with age, modeled by linear regression with an age-by-sex interaction term. Left: Adult (5-month) vs Young (1-month). Right: Aged (21-month) vs Adult (5-month). **(B)** UpSet plot showing the overlap of significantly altered cell types across the two aging transitions and across the direction of change. Top bars: intersection size (number of cell types in each combination). Right bars: total set size for each category. **(C)** Stage-specific trajectories of mean scaled newborn ratio across Young, Adult, and Aged for the three principal patterns identified in (B). **(D)** Cell-type-resolved heatmap of newborn-ratio changes across organs. Rows are organ-specific cell types and columns are tissues. Left half: Adult vs Young. Right half: Aged vs Adult. Dot color encodes log2 fold change of newborn ratio. Dot size encodes log10(number of cells). Filled circles mark FDR-significant changes (FDR < 0.05); open circles mark a significant age-by-sex interaction. Side bar plots show, per row, the number of tissues with a significant change. **(E)** Concordance between fold changes in newborn ratio (x-axis, log2FC) and global cell-type ratio (y-axis, log2FC) at the main cell-type level. Left: Young to Adult. Right: Adult to Aged. Each point is one organ-specific cell type, colored by direction of change in newborn ratio. Solid line: linear fit. Reported R² and p-value summarize the proportion of variance in global change explained by newborn change. **(F)** Pie charts cross-classifying main cell types by global-population direction (down vs up) and by newborn-ratio direction (down, no significant change, up). Left: Adult vs Young. Right: Aged vs Adult. **(G)** Same as (E), but at the cell-subtype resolution. **(H)** Same as (F), but at the cell-subtype resolution.

Most altered cell types (192 of 227) changed significantly in only one of the two transitions; the largest single group consisted of 105 cell types with reduced cell genesis from young to adult (**Figure 4B-C**). These populations spanned structural and immune lineages: endothelial cells (across 14 tissues), stromal lineages such as fibroblasts, pericytes, smooth muscle cells, and mesothelial cells, as well as B cells and innate lymphoid cells. The decline in structural lineages likely reflects the cessation of active tissue growth in early life; the reduction in lymphoid cell genesis is consistent with the contraction of primary lymphopoiesis as the peripheral immune pool becomes established(*54*, *55*), supported by the reduced EdU labeling index we observed in the spleen (**Figure 1B**). In contrast, granulocytes (neutrophils, basophils, monocytes) and plasma cells showed increased cell genesis over this interval, reflecting the buildup of innate and humoral immune capacity in early adult life **(Figure 4D**).

In contrast to the young-to-adult transition, where structural and lymphoid declines dominated, the adult-to-aged transition was characterized primarily by immune cell expansion: 48 of the 76 altered cell types showed increased cell genesis. These included adaptive lymphoid lineages (T cells, B cells, plasma cells) and myeloid populations (macrophages and monocytes) across multiple tissues (**Figure 4D**). A smaller subset (28 cell types) continued to show reduced cell genesis, mostly among structural lineages such as fibroblasts and pericytes. This pattern of elevated immune genesis alongside diminished structural renewal is consistent with the well-documented immune cell accumulation in aging tissues, a feature often termed “inflammaging” (*56*).

These broad shifts in cell genesis raised the question of how much they contribute to the parallel changes observed in the global cell population during aging. Across both transitions, fold changes in newborn ratio correlated positively with fold changes in global cell-type ratio (R² = 22.5% for young-to-adult and R² = 15.2% for adult-to-aged; **Figure 4E**). At the main cell-type level, 65.7% of cell types showed concordant directionality between newborn and global populations from young to adult, and 75.4% from adult to aged (**Figure 4F**). At the subtype level, both the correlation strength and the proportion of concordant cell types increased further (R² = 32.9% for young-to-adult and R² = 38.4% for adult-to-aged; **Figure 4G-H**). Together, these findings indicate that altered cell genesis contributes substantially to age-associated shifts in cell-population composition, with the contribution most evident at higher cellular resolution.

### Cell-renewal changes underlying aging-associated population shifts

Aging-associated changes in cell-type-specific newborn ratios could, in principle, arise from two distinct mechanisms: an intrinsic decline in cell renewal, or a shift in the global cell composition that passively redistributes the newborn fraction across cell types. To distinguish between these possibilities, we estimated a cell-renewal rate for each main cell type in each sample by dividing its newborn ratio to its global ratio. For each aging-associated cell type, we then examined two quantities: the renewal-rate baseline at the younger stage, and the fold change in renewal rate at each age transition.

Examining the young-to-adult transition first, aging-depleted cell types showed a significantly lower renewal-rate baseline than stable cell types (**Figure 5A**). This already-reduced renewal rate declined further from young to adult (**Figure 5B, left**). Expanding cell types, by contrast, started from a higher renewal-rate baseline at the young stage (Figure 5A). Their renewal rate did not change significantly from young to adult (**Figure 5B, right**). The same pattern held from adult to aged. Declining cell types showed a lower renewal-rate baseline at the adult stage (**Figure 5C**) and a further decline from adult to aged (**Figure 5D, left**), while expanding cell types again showed no significant change in renewal (**Figure 5D, right**). These contrasting patterns indicate that aging-associated expansion is largely driven by global compositional shifts, while aging-associated depletion reflects a cell-intrinsic disruption of renewal.

**Figure 5.**
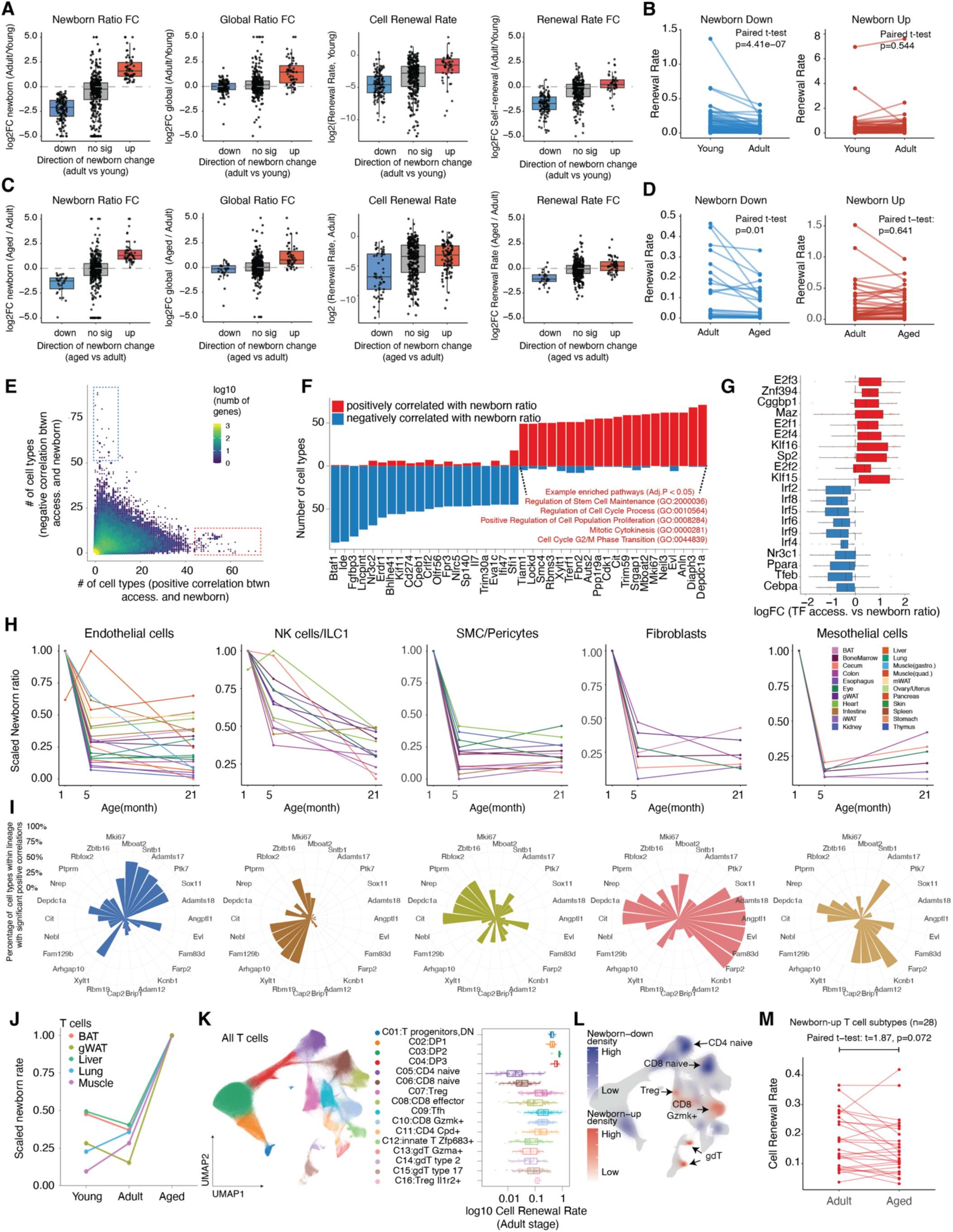
Cell-renewal dynamics and molecular features underlying newborn changes in aging. **(A)** Box plots comparing aging-associated main cell types in the young-to-adult transition, grouped by direction of newborn-ratio change. From left to right: log2 fold change of newborn ratio, log2 fold change of global ratio, baseline cell-renewal rate at the young stage (newborn ratio / global ratio), and log2 fold change of renewal rate. Each point is one organ-specific main cell type. **(B)** Paired comparison of cell-renewal rate at the young and adult stages for cell types whose newborn ratio decreases or increases from young to adult. Each line connects one organ-specific cell type across the two stages. P-values from paired t-test. **(C)** Same panel layout as (A), but for the adult-to-aged transition. Aging-depleted cell types again show a lower renewal baseline at the adult stage and a further reduction into old age, whereas aging-expanding cell types show no significant change in renewal capacity. **(D)** Same layout as (B), but for the adult-to-aged transition. P-values from paired t-test. **(E)** Scatter plot summarizing differential gene-accessibility analysis performed within each main cell type using newborn rate as a covariate. Each point is one gene; axes indicate the number of cell types in which the gene is positively (x-axis) or negatively (y-axis) correlated with the newborn ratio. Color encodes log10(number of genes) at each coordinate (point density). **(F)** Top genes recurrently correlated with newborn ratio across cell types. Bars show the number of cell types in which each gene is significantly positively (red) or negatively (blue) correlated with the newborn ratio. Selected pathway annotations from gene-set enrichment of the positively correlated genes are highlighted. **(G)** Boxplots showing the logFC of top transcription-factor motifs whose chromatin accessibility is correlated with newborn ratio across cell types. **(H)** Tissue-resolved trajectories of scaled newborn ratio across age for five broadly distributed lineages with declining newborn rates. Each line corresponds to one tissue-specific cell type. **(I)** Lineage-specific molecular features associated with the newborn ratio for each of the lineages in (H). Barplots show the proportion of tissues in which a given gene is significantly positively correlated with the newborn ratio within that lineage. **(J)** Scaled newborn ratio of T cells across young, adult, and aged stages, plotted per tissue. **(K)** UMAP embedding of all T cells pooled across tissues, colored by subtype. Right panel: distribution of log10(cell-renewal rate) at the adult stage for each subtype. **(L)** Same UMAP as in (K), highlighting cell subtypes with newborn-ratio decline and increase during the adult-to-aged transition. The aging-expanding newborn T cells map predominantly onto proliferation-active subtypes. **(M)** Lineplot showing the cell-renewal rates for aging-expanding newborn subtypes of T cells between the adult and aged stages. P-values from paired t-test.

We next asked whether the population decline of these cell types is accompanied by molecular signatures of reduced renewal. For each main cell type, we identified genes whose chromatin accessibility correlates with newborn ratio (**Figure 5E**). Across cell types, genes recurrently correlated with newborn ratio were enriched for cell cycle, cell proliferation, and stem-cell maintenance pathways (**Figure 5F**). A parallel analysis of TF motif accessibility identified E2F-family motifs (*57*, *58*) as positively correlated with newborn ratio across cell types, whereas IRF-family (interferon-response) motifs were anti-correlated (**Figure 5G**), consistent with elevated interferon signaling in aged tissues (*25*).

Beyond these shared molecular features, we examined broadly distributed cell lineages with declining newborn ratios across the body to ask whether their decline reflects conserved sets of molecular signatures (**Figure 5H–I**). Some features were positively correlated with newborn ratio broadly, including the canonical proliferation marker *Mki67* (*59*). Other features were lineage-specific yet tissue-conserved, and several have established roles consistent with their associations: *Ptk7*, a regulator of endothelial sprouting and angiogenesis (*60*), in endothelial cells; *Zbtb16*, a master regulator of innate-lymphocyte development (*61*), in innate lymphoid cells; and *Cit*, encoding the cytokinesis kinase citron (*62*), in mural cells. Together, these signatures point to lineage-specific but tissue-conserved molecular programs associated with newborn-cell genesis, and nominate candidate molecular handles for sustaining renewal in aging-depleted lineages.

We then asked what drives the expansion of cell types with increased newborn ratios in aging. T cells offered a representative case: their newborn ratio expanded with age across multiple tissues (**Figure 5J)**. Pooling T cells across the body, we resolved 16 subtypes with markedly different baseline renewal rates at the adult stage (**Figure 5K, 5L**). Thymic immature T cells showed the highest renewal rates, followed by *Gzmk+* T cells and regulatory T cells (Tregs), while naive T cells showed the lowest. During aging, newborn T cells shifted away from naive quiescent subtypes toward fast-cycling states including CD8 *Gzmk+* T cells and Tregs (**Figure 5L**). Despite this expansion, these aging-expanding subtypes themselves showed no significant change in their cell-renewal rate. (**Figure 5M**). These T-cell findings, together with the renewal decline observed in aging-depleted lineages, indicate that aging-associated changes in cell genesis arise from at least two distinct mechanisms: cell-intrinsic disruption of renewal in depleted lineages, and cell-state transitions toward proliferation-active subtypes in expanding ones. These mechanisms suggest distinct intervention strategies for sustaining tissue renewal in aging.

## Discussion

Single-cell atlases have made aging visible as a change in cellular composition. What they usually cannot show is how that composition came to be. Abundance is the endpoint of several dynamic processes, including production, differentiation, persistence, redistribution, and clearance. By coupling *in vivo* EdU labeling with single-nucleus chromatin profiling across the mouse organism, *TrackerSci* measures cell genesis within a defined labeling window and places that signal onto chromatin-defined cell states. This adds a dynamic dimension to organismal aging: tissue remodeling involves both a change in which cells are present at sampling and a change in the routes by which those cells are produced, maintained, and lost.

The first implication is that broad cell-type labels often mask the renewal-active states within each cell type. Across many organ-specific cell types, EdU-labeled nuclei mapped to molecularly distinct positions relative to their matched global populations. Among 330 main cell types with sufficient representation, 248 showed significant newborn-versus-global distribution shifts. This indicates that recent cell genesis is frequently concentrated in particular newborn-enriched states rather than spread uniformly across a parent cell type. Some states recapitulated known renewal biology, including zone-2 hepatocytes. Other states have not been previously characterized as renewal-active. These candidate progenitor-like or proliferation-active states appeared in endocrine, stromal, ovarian, epididymal, immune, and endothelial compartments. Within the endothelial compartment, the Kit+/Meox1+ newborn state appeared recurrently across tissues, suggesting that chromatin programs associated with cell production may be reused across vascular beds. Thus, *TrackerSci* exposes a layer of productive heterogeneity that static cell-type maps largely compress.

Aging also changed where newborn cells appeared along differentiation trajectories. Selected trajectories showed either accumulation of newborn cells in intermediate regions or shifts toward abnormal states, arguing against a simple model in which aging acts only by uniformly reducing proliferation or delaying differentiation. In the kidney, for example, the PEC-to-PTS3T2 trajectory revealed an age-emergent *Ror2+/Dock2+* newborn PEC-like population, illustrating how *TrackerSci* can nominate putative source routes where aging-expanding cellular states come from. In the proximal intestinal epithelial trajectory, by contrast, aging produced a stalled-differentiation pattern, with newborn cells accumulating at intermediate stages and depleting at both differentiation ends. External single-cell RNA-seq data recapitulated this signature, providing additional support for the inferred trajectory remodeling. Together with concordant chromatin and RNA programs and shifts in TF motif accessibility, these results support a model in which aging changes how newborn cells progress along differentiation trajectories. Direct fate tracing, pulse-chase experiments, and perturbation are needed to test whether these inferred bottlenecks or source-state relationships reflect true differentiation arrest or lineage transitions.

The relationship between newborn-cell dynamics and matched global populations further shows why compositional atlases are incomplete on their own. Across age transitions, changes in newborn ratio were positively associated with global cell-type composition changes, explaining 22.5% of variance from young to adult and 15.2% from adult to aged at the main-cell-type level. The association was stronger at subtype resolution (R² = 0.329 and 0.384, respectively). These percentages indicate that altered cell genesis contributes meaningfully but partially to compositional change during aging. They imply that survival, death, migration, residence time, and state transition account for additional components of compositional change. In this sense, *TrackerSci* does not turn abundance into a complete mechanistic explanation, but it separates one directly measured dynamic process from the others that must still be measured.

This decomposition clarifies the different routes by which cell populations decline or expand during aging. Cell types depleted during aging generally exhibited lower baseline renewal exacerbated by further age-related decline, combining to cause population loss through insufficient replacement. Their associated chromatin features included cell cycle, proliferation, stem-cell maintenance, E2F-family motif accessibility, and lineage-specific renewal-associated regulators, nominating programs that may accompany reduced cellular production. Expanded populations followed a different pattern. They did not generally show increased renewal rate. The T-cell compartment suggested expansion through shifts from naive states toward proliferation-active CD8 Gzmk+ and Treg subtypes. For circulating or migratory populations, EdU labeling marks newborn cells recovered from a tissue, not necessarily cells produced within that tissue. Even with this caveat, the absence of elevated renewal in expanding subtypes argues against increased local production as the dominant route. Depletion and expansion are therefore not opposite outcomes of the same aging program. They reflect distinct dynamic modes that can look similar in a static atlas.

These distinctions matter for how aging should be modeled and perturbed. A population that declines because replacement fails may require restoration of renewal capacity or niche support. A population that changes because newborn cells accumulate in intermediate states, or shift toward alternative states, may not be repaired by increasing proliferation alone. Immune expansion through state transitions or altered persistence, exemplified by the T-cell shift to CD8 Gzmk+ and Treg subtypes, would require interventions targeting these mechanisms rather than proliferation. These are not therapeutic conclusions from the present study. They are testable predictions that follow from separating production, progression, persistence, and loss.

The study also defines the limits of this dynamic view. The five-day EdU window captures recent DNA synthesis and therefore under-samples slow-cycling, quiescent and context-dependent proliferating populations. *TrackerSci* does not directly measure cell death, survival, migration, or tissue residence. Immune-cell dynamics must be interpreted in the context of systemic production and redistribution. Trajectory inference and renewal-rate decomposition are model-based. Despite these limits, *TrackerSci* adds a missing dynamic variable to aging biology. Aging-associated cellular composition is not itself a mechanism; it is the integrated outcome of how cells are produced, where they progress, how long they persist, and when they are lost.

## Acknowledgments

We thank members of the Cao lab for helpful discussions and feedback. We also thank members of the Rockefeller University High-Performance Computing Core, Comparative Bioscience Center, Genomics Resource Center, Bioinformatics Resource Center and Flow Cytometry Resource Center for their invaluable support.

## Author contributions

J.C. and W.Z. conceptualized and supervised the project; Z.L. performed all experiments; Z.L. performed all computational analyses; J.C., W.Z., and Z.L. wrote the manuscript.

## Data availability

Raw FASTQ files, processed count matrices, cell metadata, and peak metadata can be downloaded from NCBI GEO under accession number <u>GSE331488</u> and <u>GSE331489</u>.

## Code availability

Custom code and scripts used for processing sequencing reads of the EasySci-ATAC-seq library were deposited in Zenodo (*63*) (https://doi.org/10.5281/zenodo.8395492). Jupyter Notebooks for key computational analysis are available on GitHub (https://github.com/ZiyuLu041/pan-trackersci).

## Supplementary Materials

### Materials and Methods

#### Animals and organ collection

The C57BL/6 wild-type mice at one month (n=10, 5 males and 5 females), five months (n=10, 5 males and 5 females), and twenty-one months (n=12, 6 males and 6 females) were obtained from The Jackson Laboratory. Detailed information on animal individuals in this study is provided in **Table S1**. Most libraries pooled all animals to minimize batch effects between samples. The experiment IDs have been included in the metadata table of the processed dataset. Mice were housed socially and maintained on a regular 12h/12h day/night cycle. In order to analyze newborn cells, all mice were labeled with EdU (25 mg/kg, i.p. injection) at 24-hour intervals for five days. Tissues were harvested one day after the final injection. To control for circadian effects, sample harvest was performed around the same period (4-7 PM) across all individuals. Mice were euthanized utilizing inhalation of carbon dioxide (CO2). After euthanization, major organs were quickly transferred into ice-cold PBS, and the following tissues were collected: brown adipose tissue, bone marrow, cecum, colon, esophagus, eye, heart, small intestine (spanning duodenum, jejunum, and ileum), kidney, liver, lung, muscle (quadriceps and gastrocnemius), ovary and uterus, pancreas, back skin, spleen, stomach, thymus, gonadal white adipose tissue, inguinal white adipose tissue, mesenteric adipose tissue. Dissected mouse tissues were snap-frozen in liquid nitrogen and stored at -80°C. All animal procedures were in accordance with institutional regulations under the IACUC protocol 21049.

#### Nuclei extraction from multiple mammalian organs

Nuclei extraction was performed using the method from (*24*, *25*). Before extraction, frozen tissues were placed inside aluminum foil and smashed into powder on dry ice with a hammer. 10X PBS-hypotonic stock solution was prepared as follows: 6.83 g of Na2HPO4-2H2O (Sigma, 71643-250G), 3.5 g of NaH2PO4-H2O (Sigma, 71505-250G), 1.2 g of KH2PO4 (Sigma, P285-500), 1 g of KCl (Sigma, P9541-1KG), and 3 g of NaCl (Sigma, P9888-500G) in nuclease-free water to a final volume of 500 mL. On the day of nuclei extraction, 1X nuclei lysis buffer was prepared freshly as follows: final concentration of 1X PBS-hypotonic stock solution, 3 mM MgCl2 (VWR, 9CAT7062-848), 0.025% IGEPAL CA-630 (VWR, IC0219859650), 0.1% Tween-20 (Sigma, P9416-100ML), 1X cOmplete, EDTA-free Protease Inhibitor Cocktail (Sigma, 11873580001). In addition, a final concentration of 0.33M sucrose (Sigma, S0389) was included in the lysis buffer for these tissues: esophagus, stomach, intestine, cecum, colon, spleen, thymus, bone marrow, and skin. The powdered tissues were then transferred into 10-20 mL nuclei lysis buffer, followed by brief vortexing and incubation for 10-15 minutes at 4°C on a rotator. Extracted nuclei were then filtered through 40 um cell strainers (VWR, 470236-276), pelleted down at 500g for 5 minutes, resuspended in 100 - 200 uL nuclei buffer [10 mM Tris-HCl pH 7.5 (VWR, 97062-936), 10 mM NaCl (VWR, 97062-858), 3 mM MgCl2 (VWR, 97062-848), 0.025% IGEPAL CA-630 (VWR, IC0219859650), 0.1% Tween-20 (Sigma, P9416-100ML), 1X cOmplete, EDTA-free Protease Inhibitor Cocktail (Sigma, 11873580001)] for following reactions.

#### TrackerSci-ATAC library construction and sequencing

The TrackerSci-ATAC library was prepared following our prior studies (*26*). EdU staining was performed on freshly extracted nuclei using the Click-iT Plus EdU Alexa Fluor™ 647 or Alexa Fluor™ 488 Flow Cytometry assay Kit (Thermo Fisher Scientific). A 500 μL reaction buffer (prepared following the manufacturer’s protocol) supplemented with 1X cOmplete™ EDTA-free Protease Inhibitor Cocktail was added directly to the nuclei suspension, mixed well, and left at RT for 30 minutes. Then, nuclei were spun down for 5 minutes at 500g (4°C), washed once with 500 µL of 1X Click-iT saponin-based permeabilization and wash reagent, resuspended in 1 mL in nuclei buffer with 1:100 dilution of 0.25 mg/ml 4’,6-diamidino-2-phenylindole (DAPI), and FACS sorted. Alexa647 and DAPI-positive nuclei were sorted. If the number of sorted nuclei was less than one hundred thousand, the nuclei were directly distributed into 96-well plates for the Tn5 tagmentation step; otherwise, nuclei were pelleted down at 500g for 5 minutes and resuspended in nuclei buffer to a final concentration of ∼ 500 - 1000 nuclei/µL. Nuclei were mixed in a 1:1 ratio with 2X TD buffer [20 mM Tris-HCl pH 7.5, 20 mM MgCl2, 20% Dimethylformamide (Fisher, AC327175000)] and dispensed 10 µL into each well of four 96-well plates. 1 µL barcoded Tn5 was loaded into each well. Tagmentation reaction was performed at 37°C for 30 minutes with gentle shaking at 300 rpm and stopped by adding 11 µL of 2X Stop buffer [40 mM EDTA (VWR, 37062-656), 1 mM Spermidine (Sigma, S0266-1G)] to each well. Samples were pooled, washed twice, and resuspended in nuclei buffer. 5 µL of resuspended nuclei were distributed into each well of the 96-well plates. 2 µL indexed EasySci P5 ligation adapters and 3 µL ligation mix [1 µL nuclease-free water, 1 µL 10X T4 DNA ligase buffer, 1 µL T4 DNA ligase (NEB, M0202L)] were added to each well. Ligation was performed at room temperature for 30 minutes with medium-speed rocking (350g) and stopped by adding 2 µL of 18 mM EDTA to each well. After that, nuclei were pooled, washed, resuspended using nuclei buffer, and subjected to another round of FAC sort based on DAPI and Alexa647 staining to remove doublets. Then, sorted nuclei were distributed into PCR plates at 5 uL per well. Proteinase K treatment was performed by mixing each well with 0.25 μL 18.9 mg/mL proteinase K (Sigma, 3115828001), 0.25 µL 1% SDS, and 0.5 µL EB buffer, and plates were incubated at 65°C for 16 hours. Then, 2 μL 10% Tween-20 was added to each well to quench the SDS. Following on, 1 μL of 10 μM universal P5 primer (5′-AATGATACGGCGACCACCGAGATCTACAC-3′, IDT), 1 μL of 10 μM indexed P7 primer (5’-CAAGCAGAAGACGGCATACGAGAT[i7]GTGACTGGAGTTCAGACGTGTGCTCTTCCGATCT-3’, IDT), and 10 μL NEBNext High-Fidelity 2X PCR Master Mix (NEB M0541L) were added into each well. Amplification was carried out using the following program: 72°C for 5 minutes, 98°C for 30 seconds, 13-14 cycles of 98°C for 10 seconds, 66°C for 30 seconds, 72°C for 30 seconds, and a final 72°C for 5 minutes. Final PCR products were pooled and purified by column purification using a Zymo DNA Clean & Concentrator kit (Zymoresearch, D4014), followed by gel extraction using Zymoclean Gel DNA Recovery Kit (Zymoresearch, D4007) to remove adapter dimers. Library concentrations were determined by Qubit and the libraries were visualized by electrophoresis on a 2% E-Gel™ EX Agarose Gels (Invitrogen G402022).

#### Processing of sequencing reads

Base calls were converted to fastq format and demultiplexed using Illumina’s bcl2fastq/v2.19.0.316, tolerating one mismatched base in barcodes (edit distance (ED) <= 1). Then, indexed Tn5 barcodes and ligation barcodes were extracted, and corrected to their nearest barcode (edit distance (ED) <= 1). Reads with uncorrected barcodes (ED >= 2) were removed. Tn5 adaptors were removed from 5’-end and clipped from 3’-end using trim_galore/0.4.1 (https://github.com/FelixKrueger/TrimGalore). Trimmed reads were mapped to the mouse genome (mm10) using STAR/v2.5.2b with default settings. Aligned reads were filtered using samtools/v1.4.1 to retain reads mapped in proper pairs with quality score MAPQ > 30 and to keep only the primary alignment. Duplicates were removed by Picard MarkDuplicates/v2.25.2 (*64*) per PCR sample. Deduplicated bam files were converted to bedpe format using bedtools/v2.30.0 (*65*), which were further converted to offset-adjusted (+4 bp for plus strand and -5 bp for minus) fragment files (.bed). Deduplicated reads were further split into constituent cellular indices using the Tn5 and ligation barcodes, and sparse matrices counting reads overlapping with promoter regions (±1 kb around transcription start site) were generated for quality filtering. In the meantime, fragment files were used to generate h5ad files for all downstream analyses with the snap.pp.import_data() function of SnapATAC2/v2.5.1 (*66*). Cell-by-bin matrices were also generated with the SnapATAC2 function snap.pp.add_tile_matrix() containing insertion counts across genome-wide 5000-bp bins.

#### Cell filtering, dimensionality reduction, clustering, and annotations

The following analyses were performed on the dataset of each tissue separately, with EdU+ cells and global cells combined and analyzed together, as described in our prior study (*25*). After initial processing, cells with fewer than 1000 unique reads or less than 15% of reads in promoter regions were discarded. The promoter ratio cutoff was adjusted to 0.1 for the eye dataset due to the observation of a lower promoter ratio in corneal epithelial cells. Then, we used an iterative clustering strategy to detect potential doublet cells from each organ. Briefly, doublet scores were calculated for each cell using the SnapATAC2 function snap.pp.scrublet(). Meanwhile, all cells of each organ were subjected to clustering and sub-clustering analysis with spectral embedding and graph-based clustering implemented in SnapATAC2. Cells labeled as doublets (defined by a doublet probability cutoff of 0.5) or from doublet-derived sub-clusters (defined by a doublet ratio cutoff of 0.2) were manually examined and filtered out. We then generated gene activity matrices for each organ by counting the Tn5 insertions in the TSS and gene body regions for each gene using the SnapATAC2 function snap.pp.make_gene_matrix(). Gene activity matrices were then used for cell type annotations.

To identify clusters of cells corresponding to different cell types, we subjected cells after data cleaning to dimension reduction and Leiden clustering using SnapATAC2 functions snap.tl.spectral(), snap.pp.knn(), and snap.tl.leiden() with the default setting. UMAP coordinates were calculated based on the spectral embedding matrices using the function UMAP() with min_dist=0.01 from the Python package umap/v0.5.2 (*67*). For cell annotations, we first obtained a draft of annotations by integrating our chromatin data (subsampled to 2,000 cells per Leiden cluster) with the published snRNA-seq datasets (*24*) through Seurat/4.3.0.1 (*68*) label transfer, and we manually reviewed and refined the annotations for each Leiden cluster based on the accessibility of known gene markers (*25*).

#### Quantifying cell state differences between the newborn cells and the global population

To test whether newborn (EdU+) cells of a given cell type occupy a molecularly distinct molecular state from the global (DAPI) population, we computed the energy distance (*28*) between the two groups in the sub-clustering low-dimensional spectral embedding for every combination. The energy distance was defined as D = 2 * E||X-Y|| - E||X-X|| - E||Y-Y||, where the expectations are mean pairwise Euclidean distances on the spectral coordinates. Cell types with fewer than 200 cells in either group were excluded. To assess significance, we performed a balanced permutation test with downsampling to 2,000 cells per group and 1,000 random permutations. Cell types were called significantly shifted when the observed energy distance exceeded the 99.9% quantile of the null distribution pooled across all cell types (P < 0.001).

#### Trajectory analysis

To characterize cellular differentiation trajectories and their changes with aging, we focused on ten representative trajectories spanning multiple tissues and lineages, including immune cell genesis in the bone marrow (B cells, erythroid, monocytes, neutrophils, pDC) and thymus (CD4 / CD8 T cells), epithelial differentiation in the intestine, colon, and cecum, epidermal differentiation in the skin, and the parietal epithelial cell to PTS3T2 trajectory in the kidney. For each trajectory, corresponding cells were extracted and re-embedded with SnapATAC2 spectral decomposition. Pseudotime was computed on the spectral embedding using slingshot/v2.0 (*70*) with the most progenitor-like cell type set as the starting cluster. Per-cell pseudotime values were scaled to [0, 1] for downstream comparison.

To identify aging-associated cellular neighborhoods within each trajectory, we applied miloR/v1.2.0 (*41*) to the EdU+ subset of each trajectory. Per-trajectory h5ad files were converted to Milo objects, which were further built with buildGraph(k = 200, d = 30, reduced.dim = “SPECTRAL”) followed by makeNhoods(prop = 0.1, refined = TRUE). Cells were assigned to neighborhoods, and per-sample neighborhood counts were generated. Differential abundance along age was tested by fitting an edgeR (*71*) model on the neighborhood counts with age (1, 5, or 21-month) as a continuous covariate. Neighborhoods with FDR < 0.05 and |log2FC| > 0.1 were called aging-associated, and classified as aging-up (logFC > 0) or aging-down (logFC < 0). Per-trajectory summaries reported the percentage of altered neighborhoods, the distribution of altered-neighborhood pseudotime, and the logFC of significant neighborhoods.

To dissect the molecular programs underlying aging-associated changes in differentiation, we focused on the proximal intestinal epithelial cell (IEC) trajectory and integrated it with a published intestinal scRNA-seq dataset (*24*). Cells from the two modalities were aligned in a shared UMAP space, and trajectory and pseudotime labels were transferred from ATAC to RNA via KNN regression on the joint embedding. Within each modality, dynamic genes were identified by fitting a Poisson generalized additive model (pyGAM (*72*)) of normalized expression or gene-activity against pseudotime, and genes with deviance explained > 0.2 and Benjamini-Hochberg adjusted p < 0.05 were retained. Aging-associated changes were quantified by aggregating counts per sample into a pseudo-bulk matrix and applying edgeR with continuous age as the covariate. Genes with FDR < 0.05 were called aging-altered. Aging-associated genes shared between RNA and ATAC with concordant logFC direction were used for downstream interpretation. Pathway enrichment of concordant aging-down genes was performed with Enrichr (*73*) using the GO Biological Process and KEGG gene set libraries. For TF motif analysis along the same trajectory, motif accessibility per pseudo-bulk sample was quantified with motifmatchr (*74*) using JASPAR position weight matrices (*75*). The same edgeR pipeline was applied to the resulting motif-by-sample deviation matrix to identify aging-associated TF motifs.

#### Differential population analysis of newborn cells

For each main cell type or subtype, we quantified the per-sample newborn ratio as the cell-type-specific fraction within the EdU+ population, multiplied by the FACS-measured EdU+ ratio (newborn_ratio = n_EdU_cells / n_total_EdU_cells × EdU_FACS%). To capture stage-specific dynamics, we then applied regression analysis (newborn_ratio ∼ Age + Age:Sex) for the young-to-adult and adult-to-aged. The age coefficient and its t-test p-value were extracted. The model was fitted twice using each sex as the reference group, and the two outputs were combined by taking the minimum p-value and the maximum R². Samples with a total ≥ 100 cells per tissue and cell types with R² > 0.25 were retained for FDR adjustment. The age and age: sex interaction p-values were Benjamini-Hochberg corrected within each tissue. Cell types were classified as aging-up, aging-down, or interaction-significant (male-biased / female-biased) based on the FDR-corrected p-values (q < 0.05) and the sign of the age coefficient. The same procedure was applied to subtype-level counts. Fold changes of newborn ratios between age groups were also calculated and used for visualizations. For interaction-significant cell types, only the relevant sex groups were used in the fold change calculations.

#### Cell renewal analysis

To distinguish whether aging-associated changes in the newborn population are driven by cell-intrinsic renewal versus passive shifts in the global composition, we estimated a per-sample, per-cell-type renewal rate as the ratio of the newborn ratio to the global ratio. We then quantified two summary statistics for every cell type: (i) the baseline renewal rate at the younger stage and (ii) the log2 fold change in renewal rate from the younger to the older stage. Aging-depleted, aging-expanded, and stable cell types were defined from the differential newborn population analysis above, and summary statistics metrics were compared using Wilcoxon rank-sum tests. In addition, for aging-depleted and aging-expanded cells, paired t-tests were used to examine whether cell renewal rates change significantly between age groups. The same renewal-rate quantification was repeated at the subtype level.

#### Identifying newborn rate-associated changes in chromatin accessibility

To identify chromatin features whose accessibility correlates with the newborn rate of a given cell type across ages and animals, we performed differential analyses on pseudo-bulk gene-activity and peak-count matrices using edgeR/v3.36.0 (*71*). For each main cell type, single-cell counts were aggregated by mouse individual, and samples with low cell number (total cells below 10% of the per-cell-type sample mean) were excluded. Genes or peaks with TPM > 10 in at least five samples were kept for testing. Analyses were performed on cell types with at least five remaining samples and more than one unique newborn-rate value. The newborn rate per (sample, cell type) was used as a continuous covariate in the design matrix (∼ newborn_rate). DGEList objects were normalized with calcNormFactors(), dispersions were estimated with estimateDisp(), and likelihood ratio tests for the newborn-rate coefficient were performed with glmFit() followed by glmLRT(). Top tables were extracted with topTags(). Significant genes and peaks were called at an FDR of 0.05. To identify lineage-conserved molecular signatures, we counted, for each gene or peak, the number of cell types in which it was significantly positively or negatively associated with the newborn rate, and defined recurrent features as those significant in at least five cell types in the same direction. For the matched motif analysis, transcription factor motif activity was quantified per pseudo-bulk sample with chromVAR/v1.16.0 (*74*) using JASPAR and HOCOMOCO position weight matrices (*75*, *76*) on the universal peak set, and the same edgeR pipeline was applied to the resulting motif-by-sample deviation matrix to identify TF motifs whose activity correlated with the newborn rate. Pathway enrichment of newborn-rate-correlated genes was performed with Enrichr (*73*) using the GO Biological Process and KEGG gene set libraries.

## Supplementary Figures

**Figure S1.**
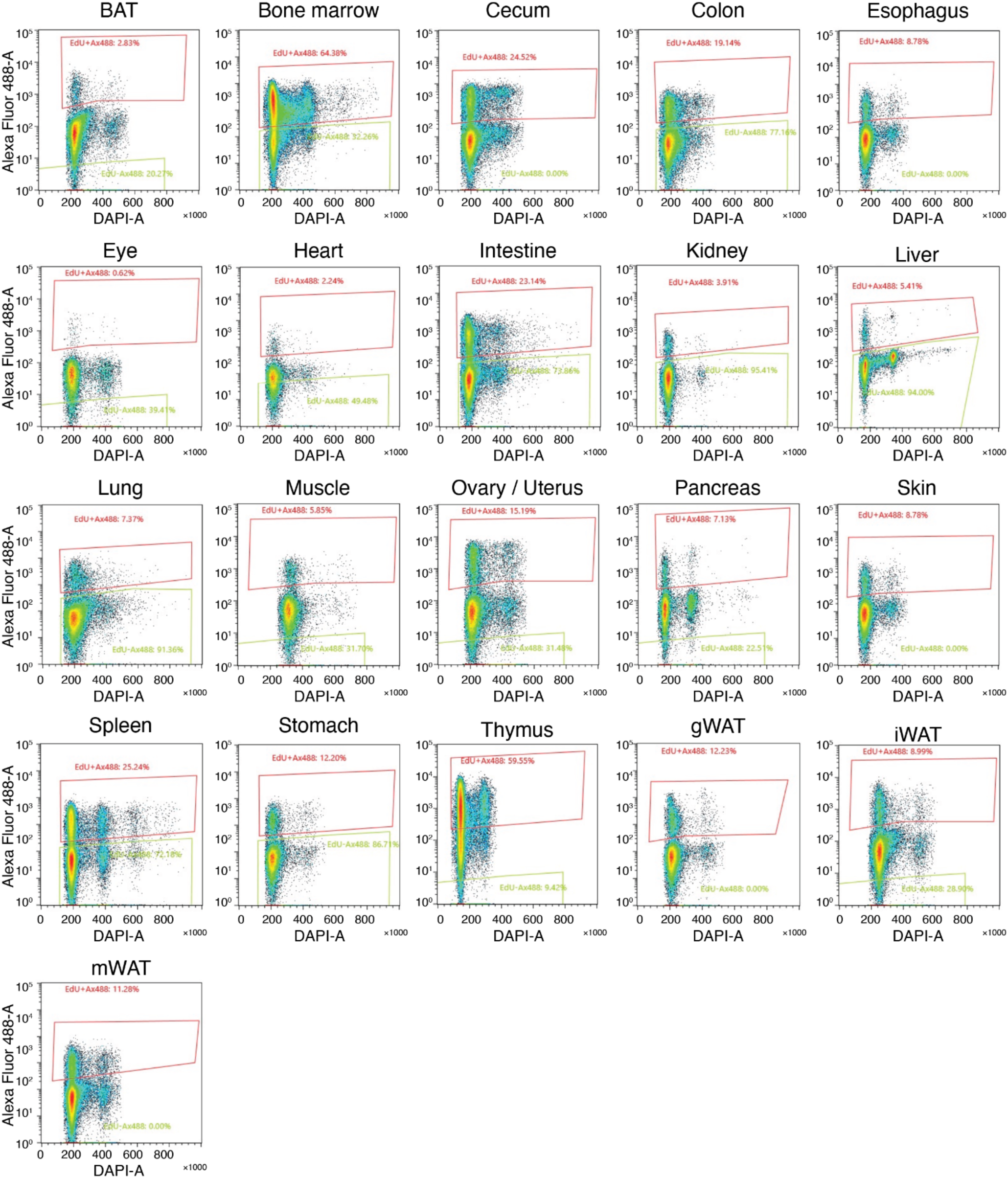
FACS enrichment of EdU+ nuclei. Representative fluorescence-activated cell sorting (FACS) scatterplots illustrating the gating strategy for EdU+ nuclei. The x-axis indicates DAPI area (linear scale), and the y-axis indicates Alexa Fluor 488 area (log scale). Red gates define the EdU+ nuclei. Data were derived from 1-month-old mice.

**Figure S2.**
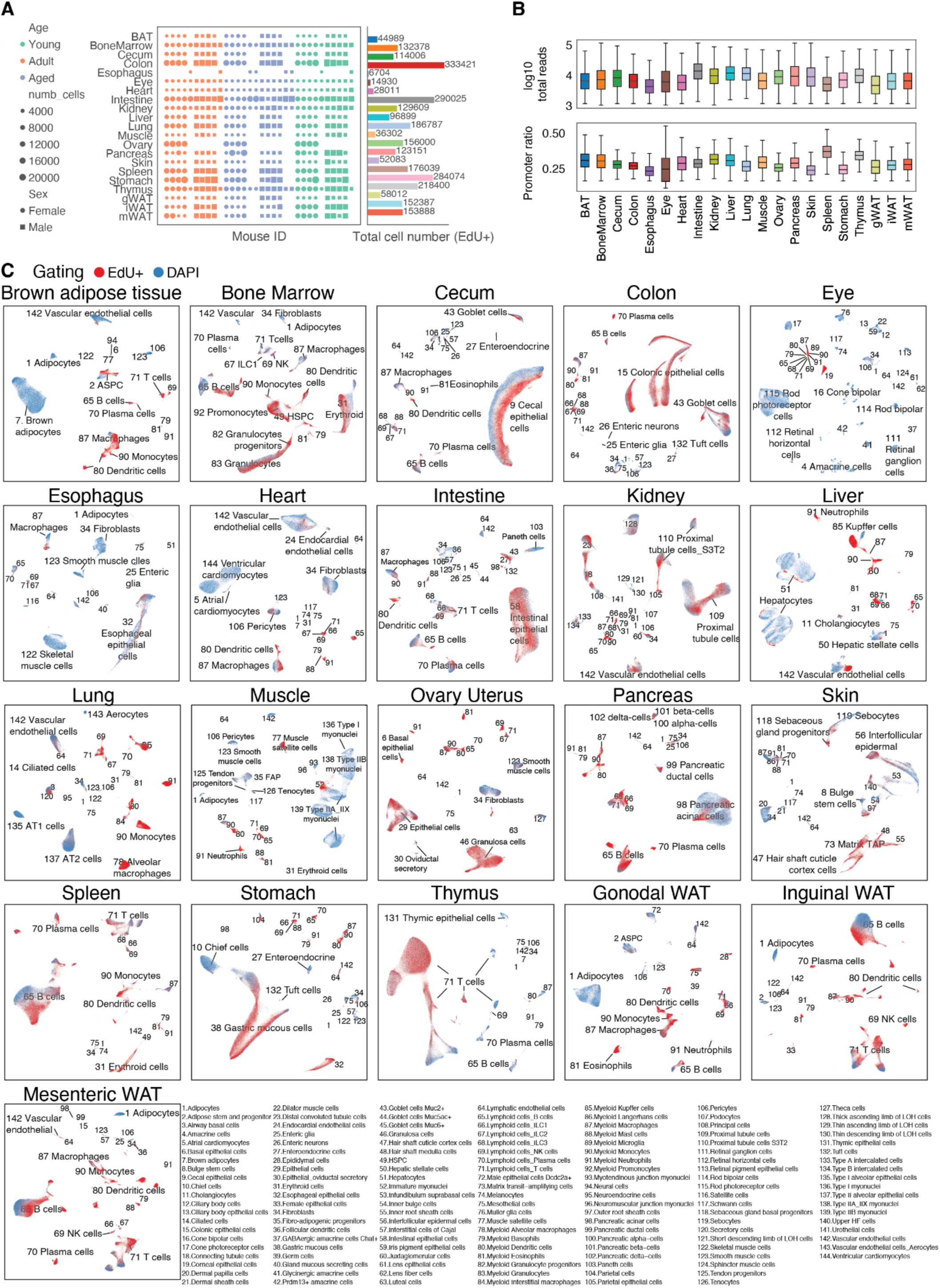
Overview of whole-organism TrackerSci-ATAC dataset. **(A)** Left: Dot plot showing the total number of EdU+ cells obtained from each individual mouse. Right: Barplot showing the total number of EdU+ cells obtained from each tissue. **(B)** Boxplot showing the log10-transformed number of unique ATAC-seq reads (top) and the ratio of reads in promoters detected per nuclei (bottom) across tissues. **(C)** UMAP visualization of all cells from each tissue, colored by EdU+ (newborn cells) or DAPI (global cells) population. Cell types were annotated and labeled as in our previous study (*25*).

**Figure S3.**
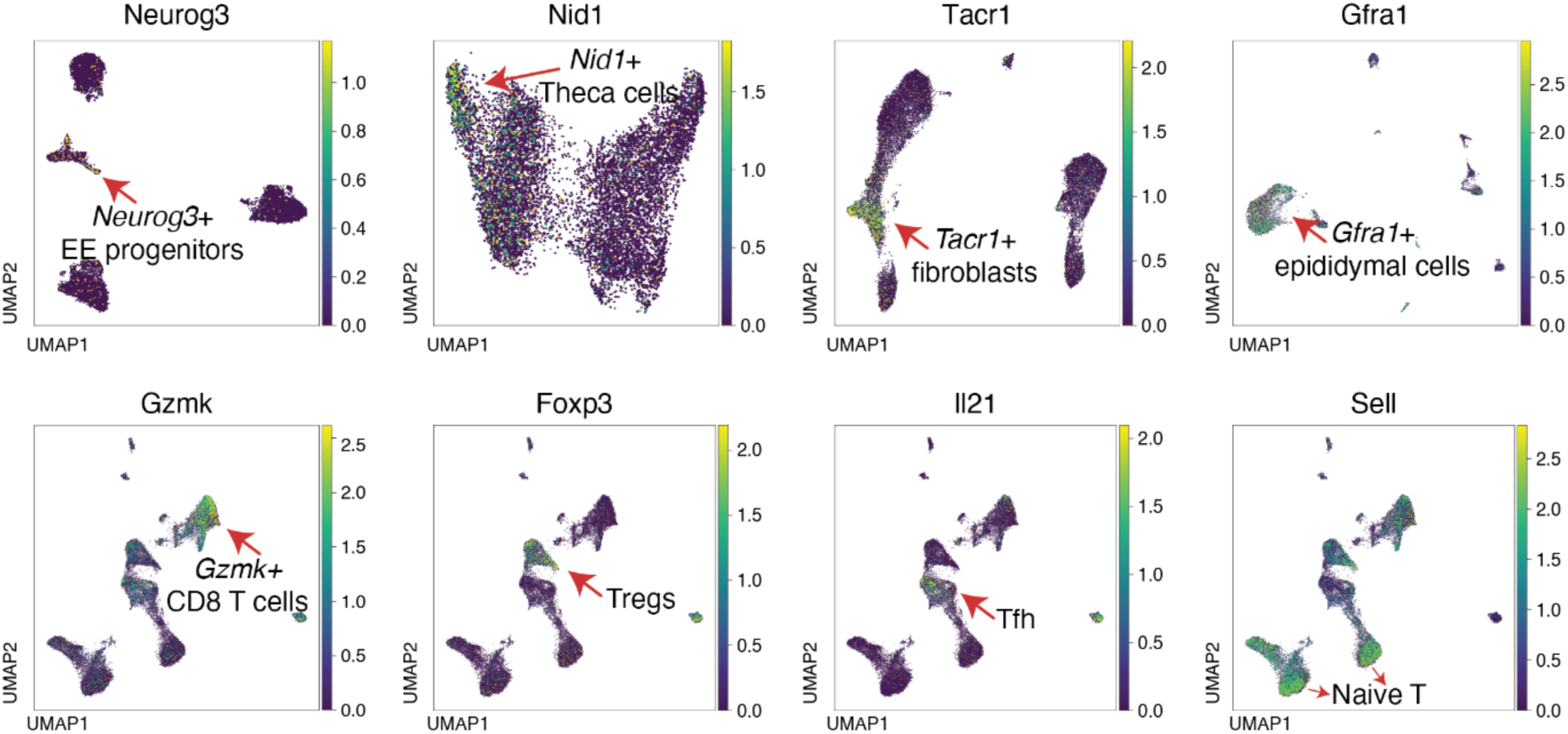
Gene activities of representative EdU-enriched subtypes. UMAP plot showing the gene marker accessibility for representative EdU+ enriched subtypes shown in Figure 2F, including *Neurog3*+ enteroendocrine (EE) progenitor cells in the stomach, *Nid1*+ theca cells in the ovary, *Tacr1*+ fibroblasts in the stomach, *Gfra1*+ epididymal cells in the gonadal white adipose tissue, and different T cell subtypes within T cell population in the Pancreas.

**Figure S4.**
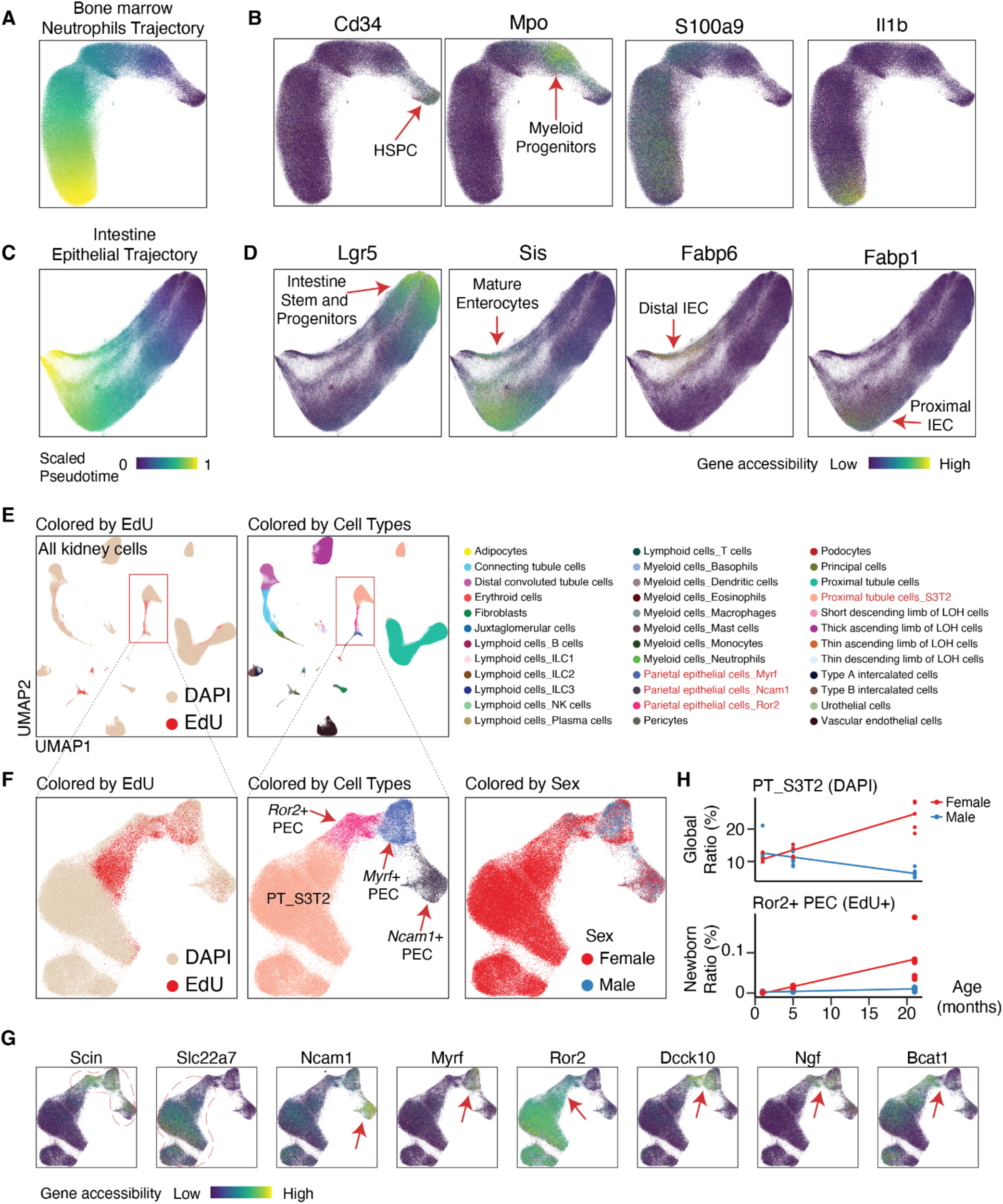
Identify cell differentiation trajectory and aging-associated changes. **(A)** UMAP visualization of the neutrophil trajectory in the bone marrow, colored by pseudotime. **(B)** UMAP plot same as (A), colored by gene accessibility of stage-specific markers, including Cd34 (HSPCs), Mpo (promyelocytes), and S100a9 and Il1b (mature neutrophils). **(C)** UMAP visualization of the intestinal epithelial trajectory in the small intestine, colored by pseudotime. **(D)** UMAP plot same as (C), colored by gene accessibility of stage-specific markers, including Lgr5 (intestinal stem cells), Sis (differentiated enterocytes), and region-specific markers for the proximal (Fabp1) and distal (Fabp6) intestine. **(E)** UMAP plot of all kidney cells, colored by EdU+ or DAPI populations. **(F)** UMAP plot of all kidney cells, colored by major cell types. **(G)** UMAP visualization of the parietal epithelial cell (PEC) trajectory in the kidney, colored by EdU/DAPI status (left), cell type (middle), and sex (right). **(H)** Chromatin accessibility of markers characterizing the transition from PECs to proximal tubule S3T2 (PT_S3T2) cells. **(I)** Dot plot showing the global fraction of PT_S3T2 cells within the DAPI population and the ratio of newborn Ror2+ PECs within the EdU+ population. Both cell types exhibit female-specific expansion.

## Supplementary Tables

**Table S1.** Metadata for the animals included in this study.

**Table S2.** Fraction of EdU-positive cells and the total number of sorted nuclei per sample across tissues.

**Table S3.** Vascular endothelial cell (VEC)-specific progenitor genes. These genes exhibited increased chromatin accessibility in the EdU-positive population compared to the DAPI-only population, uniquely within VECs.

**Table S4.** Differentially accessible and expressed genes in intestinal epithelial cells (IECs) during aging. These genes showed concordant, significant age-related changes across both scRNA-seq and scATAC-seq data. The Is_dynamic_genes column indicates whether these genes are dynamic along the IEC differentiation pseudotime.

**Table S5.** Differential abundance analysis of newborn cell populations for each cell type across tissues, comparing young and adult stages.

**Table S6.** Differential abundance analysis of newborn cell populations for each cell type across tissues, comparing adult and aged stages.

